# The *BRCA1* isoform, BRCA1-IRIS, operates independently of the full-length BRCA1 in the Fanconi anemia pathway

**DOI:** 10.1101/2022.11.02.514969

**Authors:** Andrew G. Li, Brenda C. Chan, Elizabeth C. Murphy, Ye He, Muhammed Ors, Qing Kong, Sharon B. Cantor, Joan S. Brugge, Myles Brown, David M. Livingston

## Abstract

The tumor suppressor *BRCA1* encodes multiple protein products including the canonical BRCA1-p220 (p220), which plays important roles in repair of diverse types of DNA damage. However, contributions of other *BRCA1*-encoded protein isoforms to DNA damage repair are less clear. Here, we report that the BRCA1-IRIS (IRIS) isoform has critical functions in the Fanconi anemia (FA) pathway and in repair of DNA interstrand crosslinks (ICLs). Loss of IRIS expression sensitizes cells to ICLs and impairs ICL repair. ICL formation stimulates association of IRIS with both FANCD2 and the FA core complex, which promotes FANCD2 recruitment to damage sites. The unique, *BRCA1* intron 11-encoded C-terminal tail of IRIS is required for complex formation with FANCD2 and for ICL-inducible FANCD2 mono-ubiquitylation. Collectively, our findings reveal that IRIS plays an essential role, upstream of the p220-directed HR, in the FA pathway through a previously unrecognized mechanism that depends on the IRIS-FANCA-FANCD2 interaction.

**Highlights:**

- *BRCA1* splicing isoform BRCA1-IRIS is required for interstrand crosslink (ICL) repair.
- BRCA1-IRIS interacts with FANCD2 and promotes its recruitment to sites of ICL damage.
- BRCA1-IRIS, but not BRCA1-p220, promotes ICL-inducible FANCD2 mono-ubiquitylation.
- The unique C-terminal tail of BRCA1-IRIS is essential for its function in ICL repair.

## Introduction

The canonical full-length protein product of the breast cancer susceptibility gene *BRCA1*, BRCA1-p220 (p220), plays important roles in the repair of multiple types of DNA damage, including double-strand breaks (DSBs) and interstrand crosslinks (ICLs)^1-3^. p220-directed homologous recombination (HR) is a major pathway for DSB repair and part of the Fanconi anemia (FA) pathway for ICL repair^1,2,4^. The FA pathway, consisting of 22 currently known FANC proteins and several FA-associated proteins, orchestrates the detection and removal of ICLs^5-8^. Upon recognizing ICL lesions, the FA core complex is responsible for mono-ubiquitylating the FANCD2-FANCI complex^9-11^. FANCD2 mono-ubiquitylation and recruitment to sites of ICL damage are considered essential steps for downstream processing of ICL lesions, which generates DSBs^5,12^. The resulting DSBs are resolved by HR in which p220 and other HR proteins, such as BRCA2 and RAD51, are required^4,13^. In addition to participating in HR in the FA pathway, p220 has been shown to unload the CMG helicase from stalled replication forks (SRFs) at ICL sites^14^ and to assist recruitment of FANCD2 to sites of ICL damage^3,9^.

p220 has been characterized as one of the 22 FANC proteins, termed FANCS, in the FA pathway^15^. However, prior genetic rescue studies showed that reconstitution of p220 expression, although largely restored HR, was not sufficient for a full rescue of ICL-induced defects in *Brca1*-knockout cells^16,17^, suggesting that other *Brca1* products may also be required for the response and repair of ICLs.

*BRCA1*-encoded polypeptide products include the alternatively spliced isoform BRCA1-IRIS (IRIS) protein^18^. The IRIS protein contains a unique, C-terminal tail that is encoded by a DNA segment in the intron 11 of the *BRCA1* gene and is therefore not present on the p220 protein. Unlike the well characterized function of p220 in multiple aspects of the DNA damage response and repair, a function of IRIS in this capacity has not been reported. Indeed, evidence suggests that it is not involved in the response to ionizing radiation (IR), as endogenous IRIS was not detected in IR-induced foci (IRIF)^18^. Regarding cellular response to ICLs, IRIS may interfere with cisplatin-induced apoptosis and overexpression of IRIS in ovarian cancer cell lines inhibited cell death after cisplatin treatment, doing so by transcriptionally inducing expression of the antiapoptotic protein Survivin^19,20^.

In the previous genetic rescue studies, both IRIS and p220 were deleted in *Brca1*-knockout cells^16,17^, which raises the question of whether IRIS could be the missing *Brca1* product contributing to ICL repair. Here, we report that IRIS directly participates in the FA pathway-mediated ICL repair to provide protection against ICL-induced cellular toxicity. IRIS functions upstream of the p220-directed HR in the FA pathway to promote recruitment of multiple FANC proteins to sites of ICL damage. Following ICL induction, IRIS associates with both FANCD2 and the FA core complex and promotes FANCD2 mono-ubiquitylation. The unique, *BRCA1* intron 11-encoded C-terminal tail of IRIS is essential for IRIS function in the FA pathway. Hence, the alternatively spliced *BRCA1* isoform, IRIS, operates in the FA pathway to repair ICLs, independent of p220.

## Results

### IRIS protects cells from ICL-induced cell death

To gain insight into functions of *BRCA1* isoforms in DNA damage response, we assessed the contributions of IRIS and p220 to cell survival following DNA damage. Immortalized human mammary epithelial (HME) cells expressing various short hairpin RNAs (shRNAs) were exposed to DNA-damaging agents and tested in clonal cell survival/growth assays. When compared with control (shLacZ/scr), cells expressing a hairpin specifically targeting the *BRCA1* intron 11-encoded sequence unique to the IRIS mRNA (shLacZ/shIRIS) were hypersensitive to ICL-inducing agents, such as cisplatin or MMC, but not to the DSB-inducing agent etoposide or to the SRF-inducing agent hydroxyurea (HU) (Figure 1A). When tested in parallel, cells expressing a specific hairpin targeting the 3’ UTR of the p220 mRNA (shp220/scr), in which IRIS synthesis was not interfered (Figure S1A, lane 2), displayed hypersensitivity to all four DNA-damaging drugs (Figure 1A). Enhanced cytotoxicity induced by ICL damage associated with loss of IRIS expression was not limited to epithelial cells, as human skin fibroblasts (PD846F) harboring truncating mutations in the *BRCA1* intron 11-encoded IRIS tail (IRIS^-/-^) also displayed hypersensitivity to cisplatin or MMC relative to parental (IRIS^+/+^) cells in survival/growth assays (Figure 1B). As was the case for the HME cells, loss of IRIS expression had no effect on survival of fibroblasts exposed to etoposide or HU (Figure S1B). The survival of the PD846F fibroblasts that retained one wild-type (wt) allele of the IRIS-encoding DNA segment (IRIS^+/-^) resembled the parental cells (Figure 1B), suggesting that IRIS haploinsufficiency does not affect cell survival from ICL damage. Taken together, our data suggest that IRIS is required for cell survival in response to a distinct set of genotoxins and therefore is functionally distinct from p220 whose role is ubiquitous in response to all tested DNA-damaging agents.

**Figure 1.**
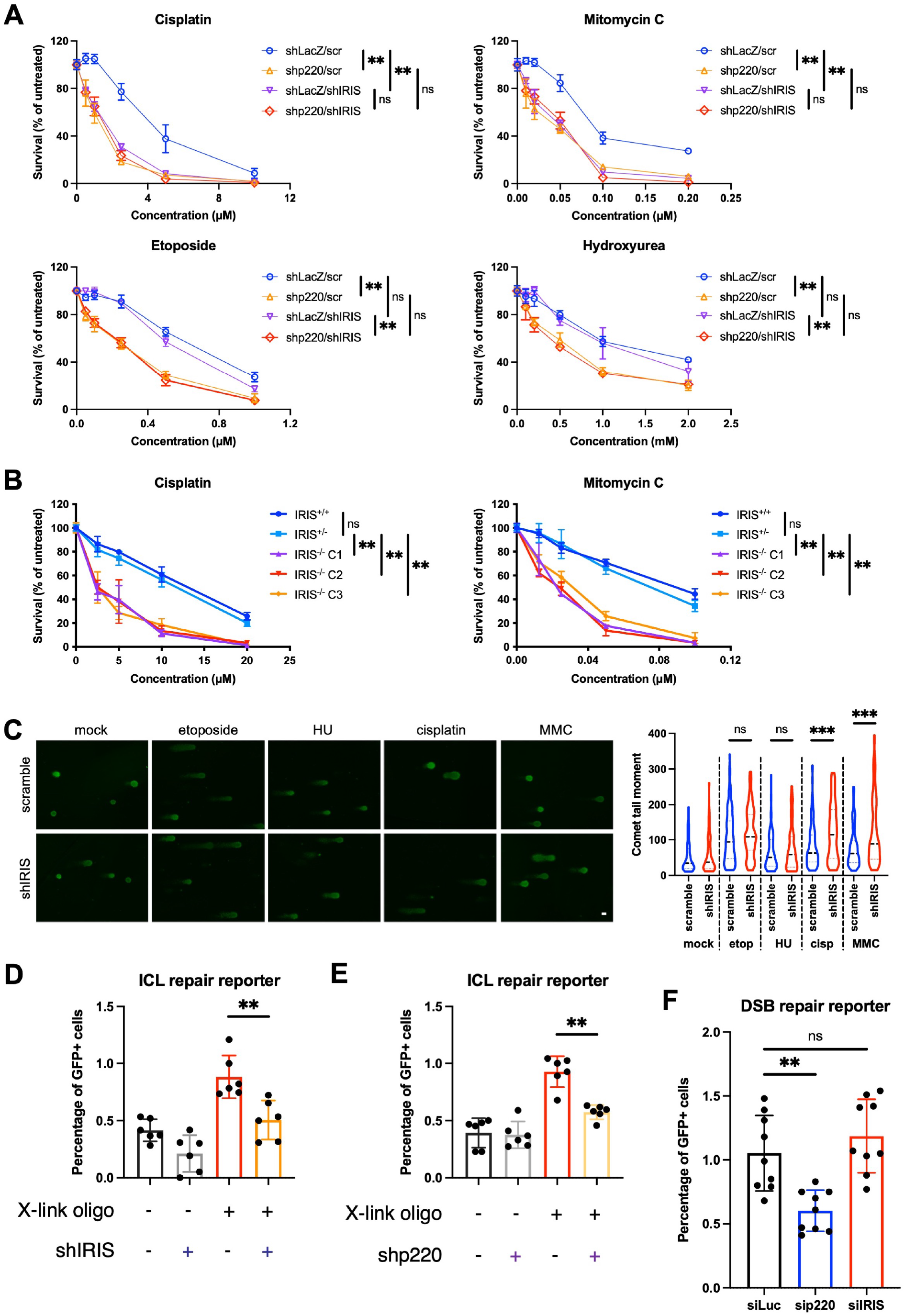
IRIS is required for ICL repair, see also Figure S1. (A) IRIS was required for HME cell survival in response to DNA interstrand crosslinkers, but not to etoposide or hydroxyurea, while p220 was required for cell survival after all four drug treatments. HME cells expressing the indicated combination of hairpins were exposed to various DNA-damaging agents for 24 hours and growth was assayed with CellTiter-Glo after a total of 5 days. Data shown are normalized mean ± SD (n = 6) of two independent experiments with triplicate wells in each experiment. scr, scramble. (B) Cisplatin and mitomycin C induced more cell death in IRIS-knockout human skin fibroblasts. Fibroblasts with the indicated IRIS genotype were treated with cisplatin or mitomycin C for 24 hours and growth was assayed with CellTiter-Glo after a total of 5 days. Data shown are normalized mean ± SD (n = 6) of two independent experiments with triplicate wells in each experiment. (C) More DNA damage, measured as comet formation, accumulated in IRIS-depleted HME cells compared with control cells after cisplatin or mitomycin C (MMC) treatment. Representative images from alkaline comet assays of cells after 24-hour treatment of each agent are shown. Quantitations of comet tail moments (n > 100) are shown on the right. etop, etoposide; HU, hydroxyurea; cisp, cisplatin. Scale bar, 20 μm. (D and E) IRIS (D) and p220 (E) were required for ICL repair in HME cells. X-link oligo, crosslinking oligonucleotide. Bars represent mean ± SD (n = 6) of two independent experiments with triplicate cultures in each experiment. (F) p220, but not IRIS, was required for HR-mediated DSB repair. Bars represent mean ± SD (n = 9) of three independent experiments with triplicate cultures in each experiment. *P* values in (A and B) were obtained using a two-way ANOVA test; *P* values in (C-F) were obtained using a Wilcoxon rank-sum test. ns, not significant; **p < 0.01; ***p < 0.001.xs

A source of ICL sensitivity following IRIS loss could be loss of p220 that functions in the FA pathway to repair ICLs^3,15^. However, this did not appear to be the case as p220 protein levels were comparable between cells devoid of IRIS expression and the corresponding control (Figures S1A and S1C, compare lane 3 to 1 in S1A). Furthermore, cells depleted of both p220 and IRIS (shp220/shIRIS) exhibited similar levels of cell death following ICL induction to cells depleted of either isoform alone (Figure 1A). The lack of additive effects in dual knockdown cells implies that p220 and IRIS function in a common pathway in response to ICL damage.

### IRIS is required for efficient cellular ICL repair

To further address the role of IRIS in the cellular response to ICL damage, we sought to determine if DNA damage accumulated in IRIS deficient cells. We found that IRIS-depleted cells exhibited increases in DNA damage as assessed by comet formation after cisplatin or MMC treatment when compared with control, while no significant changes were observed between these cells after etoposide or HU treatment (Figure 1C).

Accumulated ICLs in cells lead to replication stress^6^. Accordingly, we analyzed cellular replication stress levels by staining HME cells for phospho-CHK1, a marker of replication stress^21^. Treatment of HME cells with HU, which served as a positive control, resulted in widespread phospho-CHK1 staining in both IRIS-depleted and control cells (Figure S1D). When exposed to cisplatin or MMC, IRIS-depleted cells revealed elevated levels of phospho-CHK1 signal compared with control (Figure S1D), suggesting that more ICL lesions remained unrepaired in cells depleted of IRIS expression. Together, our data support the conclusion that IRIS is involved in ICL repair.

We next directly examined the requirement for IRIS function in ICL repair. In an ICL repair reporter assay, GFP expression scores the resolution of an ICL lesion specifically induced by a crosslinking oligonucleotide^4^. As observed in the presence of the crosslinking oligonucleotide, IRIS depletion resulted in a significant reduction in the percentage of GFP+ cells relative to control (Figure 1D). The reduction was comparable to that observed in p220-depleted cells (Figure 1E). The reduced ICL repair efficiency in IRIS-depleted cells was not a result of altered cell cycle progression, as IRIS-depleted cells showed a similar distribution of cells in each cell cycle phase when compared to control (Figure S1E). These data suggest that like p220, IRIS is required for ICL repair.

To assess a potential involvement of IRIS in the HR step of ICL repair, we analyzed the requirement for IRIS in HR-mediated DSB repair using an HR reporter^22^. We found that while p220 depletion was associated with a decrease in the level of HR repair, loss of IRIS expression did not affect the efficiency of HR repair (Figure 1F). Taken together, our data suggest that IRIS is required for ICL repair, in a step other than the p220-directed HR.

### IRIS promotes ICL-induced FANCD2 and FANCA foci

To gain insight into how IRIS regulates ICL repair, we next explored a potential functional connection between IRIS and the FA pathway. FANCD2 recruitment to sites of ICL damage is considered an essential step in the FA pathway-mediated ICL repair^5,9,12^. We therefore investigated whether IRIS regulates DNA damage-induced FANCD2 foci. Indeed, FANCD2 foci detected at sites of ICL damage were dependent on IRIS expression in both HME cells and fibroblasts (Figures 2A and 2B). FANCD2 also forms IRIF after γ-irradiation^9^. Formation of FANCD2 IRIF was not dependent on IRIS expression in HME cells (Figure S2A), suggesting that IRIS is not required for FANCD2 foci induced by DSBs.

**Figure 2.**
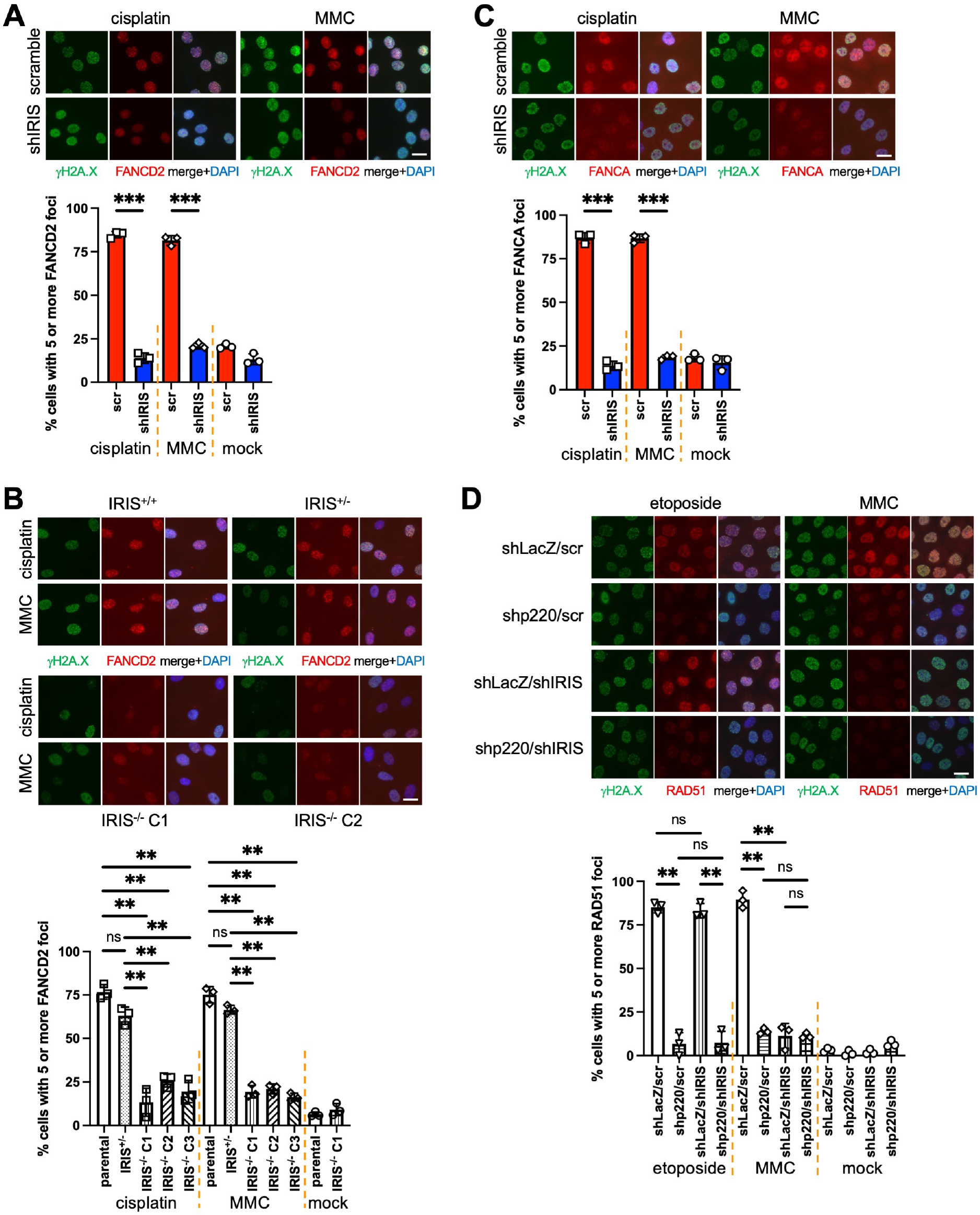
IRIS promotes ICL-induced FANCD2 and FANCA foci, see also Figure S2. (A) Formation of FANCD2 foci in control vs. IRIS-depleted HME cells exposed to cisplatin or MMC. Quantifications of percentage of FANCD2 foci-positive cells are shown on the bottom. (B) Defects in formation of FANCD2 foci in IRIS-knockout fibroblasts following cisplatin or MMC treatment. IRIS genotypes of the fibroblasts are indicated. Quantifications of percentage of FANCD2 foci-positive cells are shown on the bottom. (C) IRIS depletion resulted in diminished FANCA foci in HME cells following ICL formation. Quantifications of percentage of FANCA foci-positive cells are shown on the bottom. (D) IRIS was required for RAD51 foci after MMC treatment, whereas it was dispensable for etoposide-induced RAD51 foci. Quantifications of percentage of RAD51 foci-positive cells are shown on the bottom. Representative images of immunofluorescence of cells treated with the indicated drug for 24 hours before immunostaining using the indicated antibodies are shown. Bars represent mean ± SD (n = 3) of three independent experiments. *P* values were obtained using a two-tailed Student’s t test. ns, not significant; **p < 0.01; ***p < 0.001. Scale bar, 20 μm.

We further examined the role of IRIS in regulating ICL-induced foci of FA pathway proteins upstream of FANCD2. ICL-induced foci of FANCA, a component of the multi-subunit FA core complex that detects ICL lesions^7,8^, were dependent on IRIS expression (Figure 2C). In contrast, ICL-induced foci of UHRF1, a known ICL sensor^23,24^, were not affected by IRIS depletion in HME cells (Figure S2B). The observed defects in recruitment of FANCD2 and FANCA to sites of ICL damage associated with IRIS loss were not a consequence of decreases in their expression or depletion of S phase cells, as neither significant changes in abundance of these proteins nor reduced accumulations of S phase cells after cisplatin or MMC treatment were detected in IRIS-depleted cells (Figures S2C and S2D). Together, these results suggest that IRIS promotes ICL-inducible FANCA and FANCD2 foci and that IRIS is not involved in ICL detection.

We next investigated whether IRIS-dependent FANCD2 and FANCA foci were linked to ICL repair by examining the foci of RAD51, recently identified as FANCR and found to function as a p220-dependent recombinase in HR^25-28^. As shown in Figure 2D, RAD51 foci were reduced in IRIS-depleted cells (shLacZ/shIRIS) to a level comparable to that observed in p220-depleted cells (shp220/scr) after MMC treatment. Taken together, our data suggest that IRIS functions upstream of the p220-directed HR in the FA pathway and in a step between the initial ICL damage sensing and FANCD2 recruitment to damage sites.

### IRIS interacts with FANCD2 and FANCA in response to ICL damage

To explore the mechanism under which IRIS facilitates ICL-inducible FANCD2 and FANCA foci, we tested whether IRIS interacts with these proteins. We found that FANCD2 co-immunoprecipitated (co-IP) with FANCA, as reported previously^29^, and IRIS after cisplatin or MMC treatment (Figure 3A), whereas FANCD2 co-IPed with neither IRIS nor FANCA in etoposide-treated HME cells (Figure S3A). These data suggest that ICL damage stimulates formation of a complex containing these three proteins. Interestingly, FANCD2 failed to co-IP with FANCA in IRIS-depleted cells when tested in parallel (Figure 3A), suggesting that IRIS is responsible for the integrity of this protein complex.

**Figure 3.**
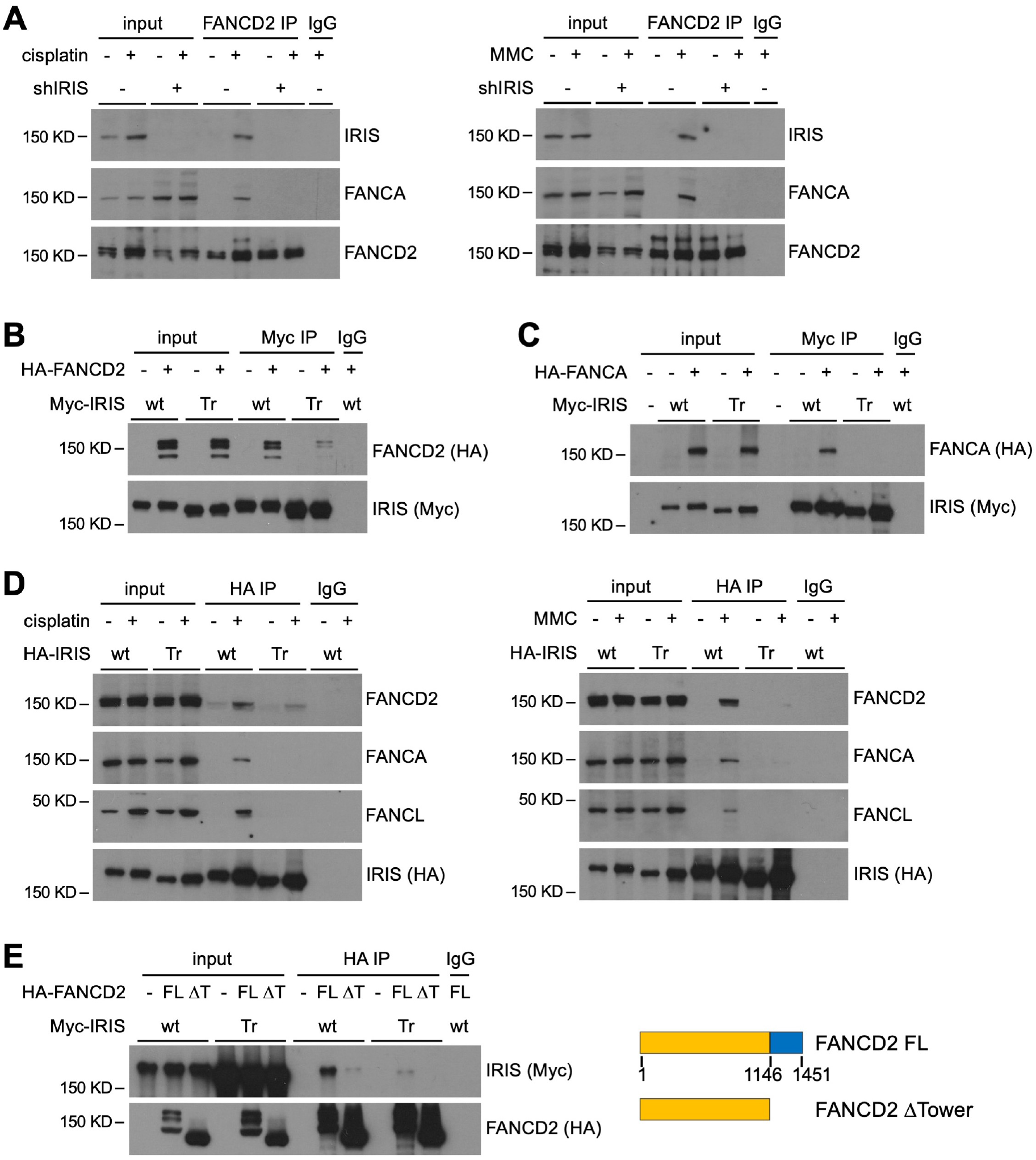
IRIS interacts with FANCD2 and FANCA in response to ICL damage, see also Figure S3. (A) IRIS associated with FANCD2 and FANCA after cisplatin or MMC treatment. Nuclear extracts (NEs) of the indicated HME cells were prepared after 24-hour drug treatment and used for immunoprecipitation (IP) with FANCD2 antibody followed by immunoblotting. (B) The *BRCA1* intron 11-encoded tail of IRIS was important for FANCD2 interaction. Whole cell extracts (WCEs) of transfected 293T cells were used for immunoprecipitation (IP) with Myc antibody, and the immune complexes were analyzed by immunoblotting. (C) wt IRIS, but not Tr IRIS, interacted with FANCA. WCEs of transfected 293T cells were used for IP with Myc antibody followed by immunoblotting. (D) Tr IRIS was unable to associate with FANCD2 and certain subunits of the FA core complex following exposure of HME cells to cisplatin or MMC. After 24-hour drug treatment, NEs from the indicated HME cells were used for IP with HA antibody followed by immunoblotting. (E) The Tower domain of FANCD2 and the C-terminal tail of IRIS were required for FANCD2-IRIS interaction. WCEs of transfected 293T cells were used for IP with HA antibody followed by immunoblotting. A schematic representation of full-length (FL) and Tower domain-deletion (ΔT, ΔTower) mutant FANCD2 proteins is shown. Numbers indicate positions of amino acid residues of the FL protein.

We next investigated whether the unique C-terminal tail of IRIS plays a role in this protein complex formation. Unlike wt IRIS, a truncated (Tr) IRIS species, which lacks the last 33 amino acid residues encoded by *BRCA1* intron 11 (Figure S3B), failed to efficiently co-IP with either FANCD2 or FANCA (Figures 3B and 3C). These results suggest that the unique C-terminal tail of IRIS is required for IRIS to bind to FANCD2 and FANCA.

As our attempt to co-IP FANC proteins using an IRIS tail-specific antibody failed, likely due to the steric epitope shielding by associated proteins, we introduced hairpin-resistant, N-terminal HA-tagged wt or Tr IRIS into endogenous IRIS-depleted HME cells. The expression levels of exogenous IRIS proteins were comparable to the endogenous levels in control cells (Figure S3C). When extracts of these cells were subjected to HA IP, wt IRIS co-IPed with FANCD2 and FANCA after cisplatin or MMC treatment (Figure 3D). In addition, wt IRIS co-IPed with FANCL, another subunit of the FA core complex^30^ (Figure 3D), suggesting that ICL formation stimulates association of IRIS with the FA core complex. In cells exposed to etoposide, wt IRIS did not co-IP with these proteins (Figure S3D). When tested in parallel, Tr IRIS failed to co-IP with these proteins (Figures 3D and S3D), suggesting that the C-terminal tail of IRIS is required for ICL-induced association of IRIS with the FA core complex.

The Tower domain of the FANCD2 protein was reported to be critical for ICL-inducible FANCD2 foci^31^. Thus, we investigated whether IRIS interacts with the Tower domain. In 293T cells, the amount of wt IRIS associated with the Tower domain-deleted FANCD2 (FANCD2 ΔTower)^31^ by co-IP was dramatically reduced compared with the full-length FANCD2 (FANCD2 FL) (Figure 3E). Furthermore, the weak association of Tr IRIS to FANCD2 FL was completely abolished when tested with FANCD2 ΔTower (Figure 3E). These data suggest that the C-terminal tail of IRIS and the Tower domain are necessary for the IRIS-FANCD2 interaction. Taken together, our data support the conclusion that induction of ICL damage stimulates association of IRIS with both FANCD2 and the FA core complex and that this requires the unique C-terminal tail of IRIS.

### The unique C-terminal tail of IRIS is required for ICL-induced FANCD2 and FANCA foci

We further examined whether the association of IRIS with FANCD2 and FANCA occurs in ICL-induced foci. We found that in endogenous IRIS-depleted HME cells expressing wt HA-IRIS, IRIS signals were detected by confocal microscopy in FANCD2- and FANCA-containing foci after cisplatin or MMC treatment (Figures 4A and 4B). The IRIS/FANCD2-overlapping foci were specifically induced by ICLs, as IRIS did not appear in etoposide-induced FANCD2 foci (Figure S4A). In addition, IRIS foci were detected partially overlapping with γH2A.X foci by confocal microscopy (Figure S4B). These data suggest that IRIS colocalizes with FANCD2 and FANCA at sites of ICL damage.

**Figure 4.**
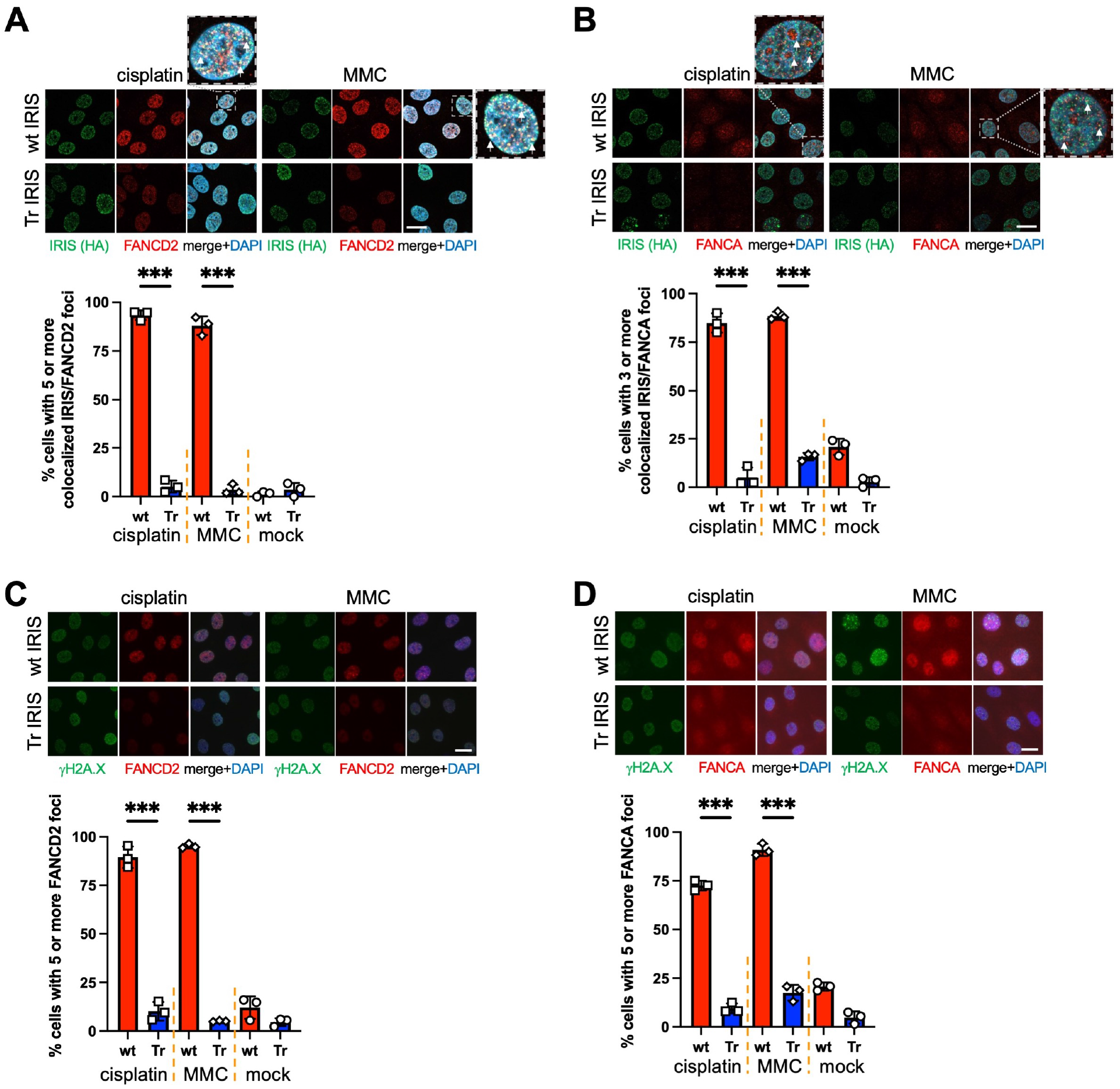
The unique C-terminal tail of IRIS is required for ICL-induced FANCD2 and FANCA foci, see also Figure S4. (A) FANCD2 colocalized with wt, but not Tr, IRIS in ICL-induced foci. Arrows point to some IRIS/FANCD2 colocalized foci. Quantifications of percentage of cells positive for IRIS/FANCD2-overlapping foci are shown on the bottom. (B) Colocalization of FANCA and wt IRIS in ICL-induced foci. Arrows point to some IRIS/FANCA colocalized foci. Quantifications of percentage of cells positive for IRIS/FANCA-overlapping foci are shown on the bottom. (C and D) Tr IRIS failed to rescue FANCD2 (C) or FANCA (D) foci in IRIS-depleted HME cells exposed to cisplatin or MMC. Quantifications of percentage of FANCD2 or FANCA foci-positive cells are shown on the bottom. Representative images of immunofluorescence of HME cells treated with the indicated drug for 24 hours before immunostaining using the indicated antibodies are shown. Zoomed-in areas containing a single nucleus each are shown next to their corresponding merged panels in (A and B). Bars represent mean ± SD (n = 3) of three independent experiments. *P* values were obtained using a two-tailed Student’s t test. ***p < 0.001. Scale bar, 20 μm.

Compared with wt IRIS-expressing cells, Tr IRIS-expressing cells exhibited reduced FANCD2 and FANCA foci in response to ICL damage (Figures 4A and 4B), suggesting that the unique C-terminal tail of IRIS is important for ICL-inducible FANCD2 and FANCA foci. To test this idea, we analyzed FANCD2 and FANCA foci with γH2A.X co-staining. Indeed, FANCD2 and FANCA localized to γH2A.X foci in response to ICL damage in cell expressing wt IRIS, but not in cells expressing Tr IRIS (Figures 4C and 4D). Neither wt nor Tr IRIS affected etoposide-induced FANCD2 foci (Figure S4C). These results suggest that the C-terminal tail of IRIS is required for ICL-induced FANCD2 and FANCA foci. Taken together, our data suggest that IRIS promotes recruitment of FANCD2 to sites of ICL damage by binding to both FANCD2 and the FA core complex at damage sites and that this function requires the C-terminal tail of IRIS.

### IRIS promotes mono-ubiquitylation of FANCD2 following ICL formation

In addition to formation of FANCD2 foci, the Tower domain of the FANCD2 protein was implicated in regulating ICL-inducible FANCD2 mono-ubiquitylation^31^. Our data showing the involvement of the Tower domain in IRIS interaction prompted us to assess the role of IRIS in this key post-translational modification on FANCD2. Western blot analyses were performed using nuclear extracts from HME cells exposed to DNA-damaging agents. As observed previously^9,10,32^ and as demonstrated here, the endogenous FANCD2 protein was ubiquitylated, as represented by the upper band on FANCD2 blots (denoted as FANCD2-L), and the FANCD2-L levels increased in control cells (shp220-/shIRIS-) after cisplatin or MMC treatment (Figure 5A, compare lane 2 to 1 in each panel). In IRIS-depleted cells (shp220-/shIRIS+), increases in FANCD2-L levels were significantly reduced following ICL formation (Figure 5A, compare lane 4 to 2 in each panel). When analyzed in parallel, ICL-induced increases in FANCD2-L levels were not affected by loss of p220 expression (shp220+/shIRIS-) (Figure 5A, compare lane 6 to 2 in each panel), as reported previously^3^. In cells treated with etoposide, IRIS depletion did not alter the induction of FANCD2 mono-ubiquitylation (Figure S5A, compare lane 4 to 2). These data suggest that IRIS is required, whereas p220 is dispensable, for ICL-inducible FANCD2 mono-ubiquitylation.

**Figure 5.**
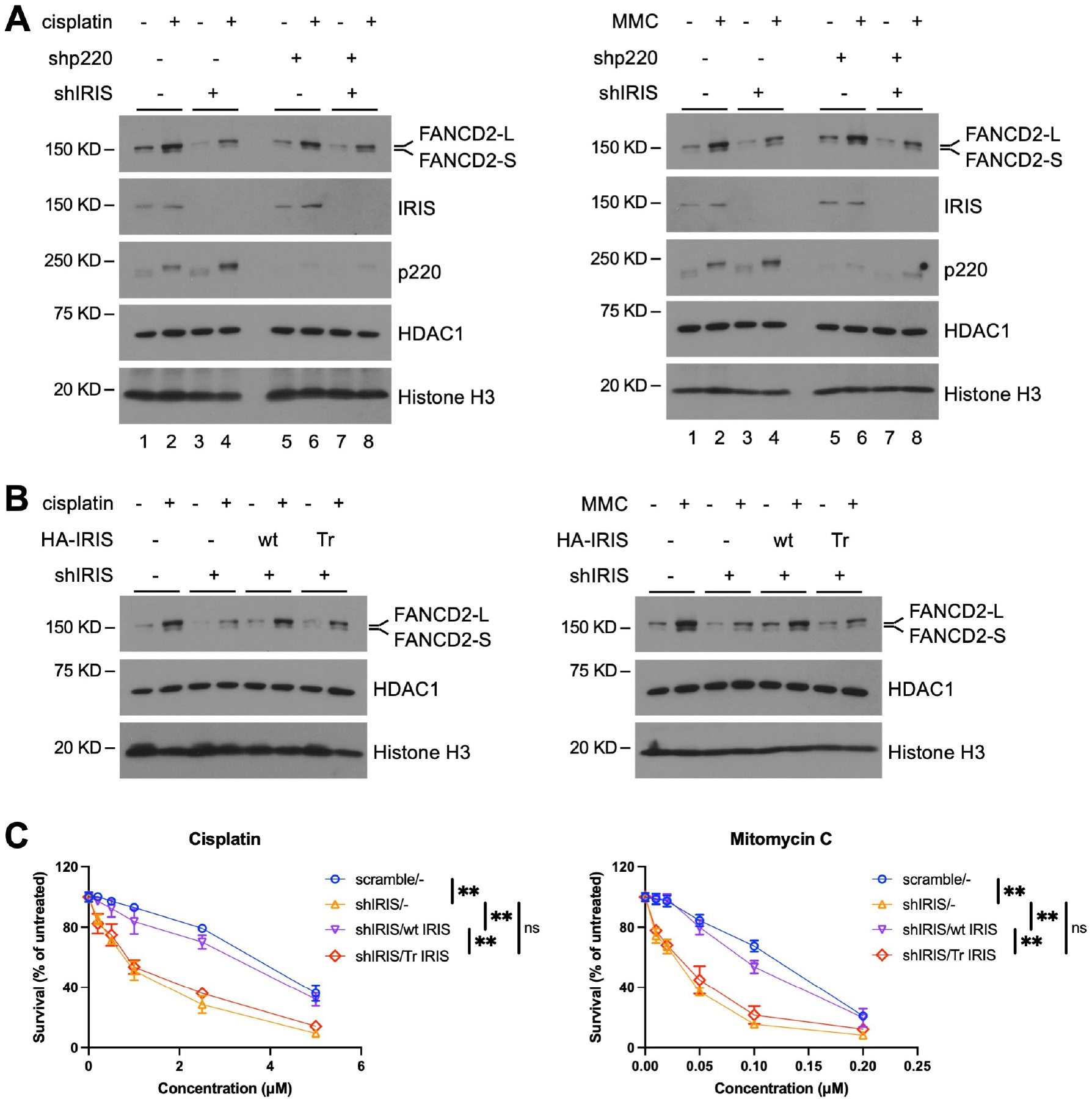
The C-terminal tail of IRIS is required for ICL-induced FANCD2 mono-ubiquitylation and cell survival from ICL damage, see also Figure S5. (A) IRIS depletion resulted in reductions in mono-ubiquitylation of FANCD2 in response to ICL damage. NEs of the indicated HME cells were prepared after 24-hour drug treatment and subjected to immunoblotting. (B) Expression of wt, but not Tr, IRIS restored FANCD2 mono-ubiquitylation in response to ICL damage. After 24-hour drug treatment, NEs from the indicated HME cells were used for Western blotting analysis. (C) Expression of wt, but not Tr, IRIS in IRIS-depleted HME cells rescued hyper toxicity induced by ICL damage. HME cells expressing the indicated hairpin and cDNA were exposed to cisplatin or mitomycin C for 24 hours and growth was assayed with CellTiter-Glo after a total of 5 days. Data shown are normalized mean ± SD (n = 6) of two independent experiments with triplicate wells in each experiment. *P* values were obtained using a two-way ANOVA test. ns, not significant; **p < 0.01.

We further examined the role of the C-terminal tail of IRIS in FANCD2 mono-ubiquitylation. Expression of wt IRIS rescued the defect in ICL-induced FANCD2 mono-ubiquitylation in IRIS-depleted HME cells, whereas expression of Tr IRIS had little effect (Figure 5B). Expression of neither wt nor Tr IRIS influenced FANCD2 mono-ubiquitylation after etoposide treatment (Figure S5B). These data suggest that the unique C-terminal tail of IRIS is important for ICL-induced FANCD2 mono-ubiquitylation.

Given the importance of the C-terminal tail of IRIS for FANCD2 mono-ubiquitylation and recruitment to sites of ICL damage in the FA pathway, we next examined its role in IRIS-dependent cell survival from ICL damage. As shown in Figure 5C, expression of wt IRIS, but not Tr IRIS, in endogenous IRIS-depleted HME cells reversed hypersensitivity to cisplatin or MMC. Neither wt nor Tr IRIS expression had an impact on cell survival after etoposide or HU treatment (Figure S5C). These data suggest that the C-terminal tail of IRIS is indispensable for IRIS to protect cells from ICL-induced cell death. Taken together, our data strongly argue that the unique C-terminal tail of IRIS is essential for IRIS function to promote FANCD2 mono-ubiquitylation and recruitment to sites of ICL damage and for the IRIS-dependent cell survival from ICL damage.

### IRIS and p220 play non-redundant roles in the FA pathway

The *BRCA1* intron 11-encoded polypeptide segment in the C-terminal tail of IRIS is not present on the p220 protein. Thus, our results suggest that IRIS and p220 function non-redundantly in the FA pathway. Indeed, our data showed that IRIS, but not p220, promotes ICL-inducible FANCD2 mono-ubiquitylation (Figure 5A). We next tested whether p220 regulates other IRIS-dependent steps in the FA pathway, such as FANCA and FANCD2 foci. We found that MMC-induced FANCA foci were not affected by p220 depletion (Figure 6A), suggesting that p220 is not required for ICL-inducible FANCA foci. In contrast, reduced FANCD2 foci were observed in p220-depleted cells after MMC treatment (Figure S6A), consistent with previous results^3,9^. This observation raised the question of whether p220 contributed to IRIS-dependent FANCD2 foci. As our data suggest that the IRIS-FANCD2 interaction is important for FANCD2 foci, we next investigated whether p220 regulates the ICL-stimulated IRIS-FANCA-FANCD2 complex. We found that following ICL formation, FANCD2 co-IPed with IRIS and FANCA in p220-depleted cells, at levels comparable to those co-IPed with FANCD2 in control cells (Figure 6B). These data suggest that p220 is not required for the IRIS-FANCA-FANCD2 protein complex in response to ICL damage. Together, these results suggest that IRIS, but not p220, promotes FANCA recruitment to sites of ICL damage and that IRIS and p220 function through distinct mechanisms to promote ICL-inducible FANCD2 foci.

**Figure 6.**
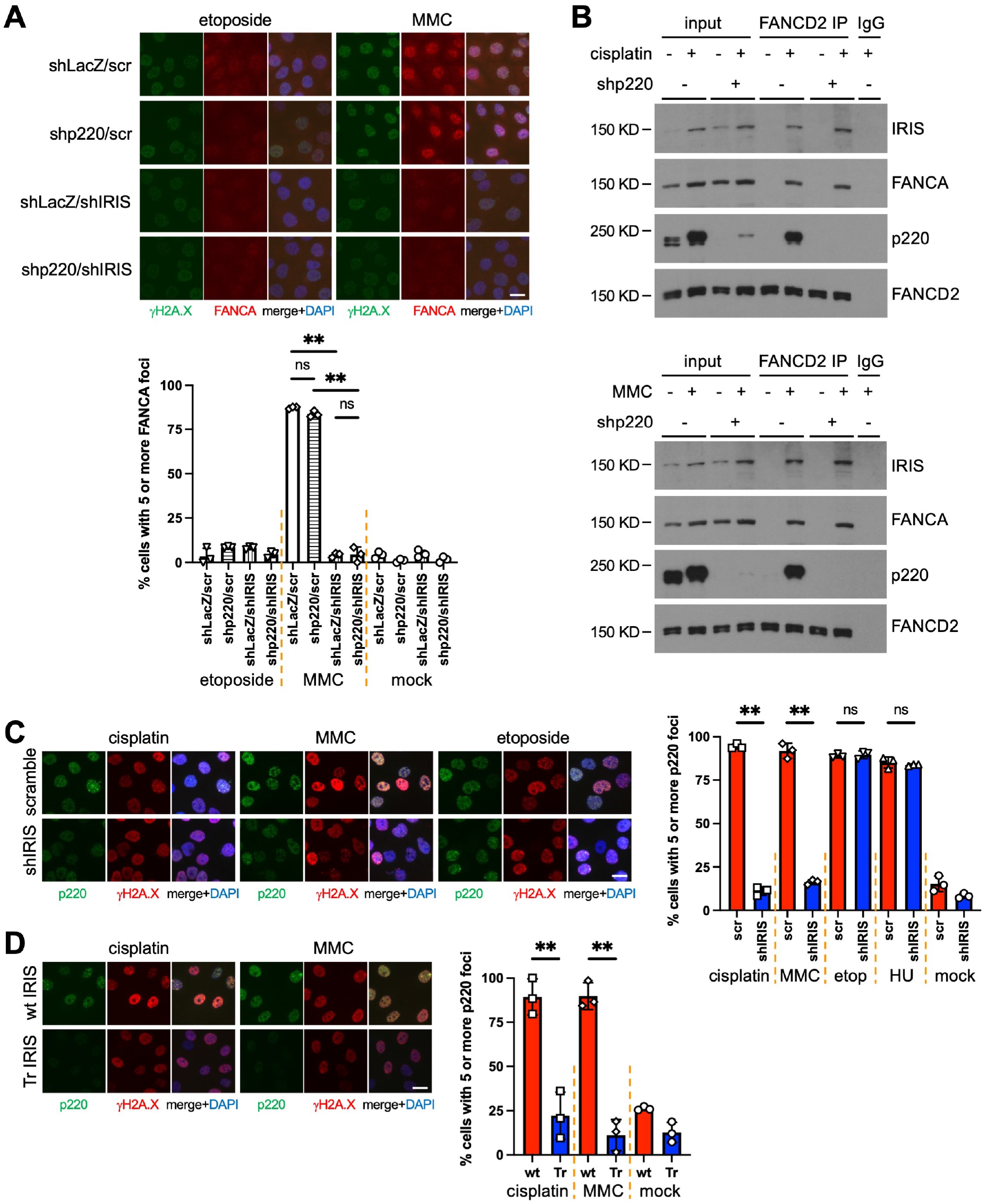
IRIS and p220 play non-redundant roles in the FA pathway, see also Figure S6. (A) IRIS, but not p220, was required for FANCA foci in HME cells after MMC treatment. Quantifications of percentage of FANCA foci-positive cells are shown on the bottom. (B) p220 was dispensable for the ICL-induced IRIS-FANCA-FANCD2 complex. NEs of the indicated HME cells were prepared after 24-hour drug treatment and used for IP with FANCD2 antibody followed by immunoblotting. (C) Loss of IRIS expression impaired p220 foci in HME cells in response to cisplatin or MMC, but not to etoposide or HU. Quantifications of percentage of cells positive for p220 foci are shown on the right. (D) Expression of wt, but not Tr, IRIS rescued ICL-inducible p220 foci in IRIS-depleted HME cells. Quantifications of percentage of p220 foci-positive cells are shown on the right. Representative images of immunofluorescence of HME cells treated with the indicated drug for 24 hours before immunostaining using the indicated antibodies are shown. Bars represent mean ± SD (n = 3) of three independent experiments. *P* values were obtained using a two-tailed Student’s t test. ns, not significant; **p < 0.01. Scale bar, 20 μm.

In addition to FANCD2 recruitment to damage sites, FANCD2 mono-ubiquitylation is critical for ICL lesion processing to generate DSBs^5,12^. The requirement for IRIS in ICL-induced FANCD2 mono-ubiquitylation (Figure 5A) suggests that IRIS functions upstream of the p220-directed HR in the FA pathway. We therefore investigated whether IRIS regulates p220 function in ICL repair. We found that IRIS depletion resulted in reduced p220 foci after cisplatin or MMC treatment, but not after etoposide or HU treatment or exposure to γ-irradiation (Figures 6C and S6B), suggesting that IRIS is required for p220 function in ICL repair, but not for p220-mediated repair of other damage types. We further examined the role of the unique C-terminal tail of IRIS in regulating p220 foci. We found that expression of Tr IRIS, when compared with wt IRIS, resulted in reduced p220 foci following ICL formation (Figure 6D). Neither wt nor Tr IRIS affected etoposide-induced p220 foci (Figure S6C). These data suggest that the unique C-terminal tail of IRIS is required for ICL-inducible p220 foci. Taken together, our data suggest that while IRIS function is independent of p220 in response and repair of ICL damage, the p220 function in the FA pathway, at least in part, is dependent on IRIS.

### IRIS and FANCA function cooperatively in response to ICL damage

To gain further insight into the p220-independent function of IRIS in the FA pathway, we generated heterozygous IRIS and/or heterozygous *FANCA* deleted as well as homozygous IRIS or *FANCA* deleted skin fibroblasts using the CRISPR-mediated gene editing. The IRIS/*FANCA* compound heterozygous cells (sgIRIS^+/-^;sg*FANCA*^+/-^ C1 and C2) revealed similar levels of cell death after cisplatin or MMC treatment compared to cells homozygously deleted for IRIS or *FANCA* (sgIRIS^-/-^ and sg*FANCA*^-/-^) as well as patient-derived *FANCA*^-/-^ cells (Figure 7A). These IRIS/*FANCA* compound heterozygous cells and the IRIS or *FANCA* null cells exhibited greater cell death after cisplatin or MMC treatment than either of the single heterozygous IRIS or *FANCA* cells (sgIRIS^+/-^ or sg*FANCA*^+/-^) or parental cells (Figure 7A). The hypersensitivity of the compound heterozygote cells to DNA interstrand crosslinkers was not a consequence of complete loss of expression of either IRIS or FANCA protein in these fibroblasts, as both proteins were detected by immunoblotting in sgIRIS^+/-^;sg*FANCA*^+/-^ cells (Figure 7B). These data suggest that IRIS cooperates with FANCA to prevent ICL-induced cell death.

**Figure 7.**
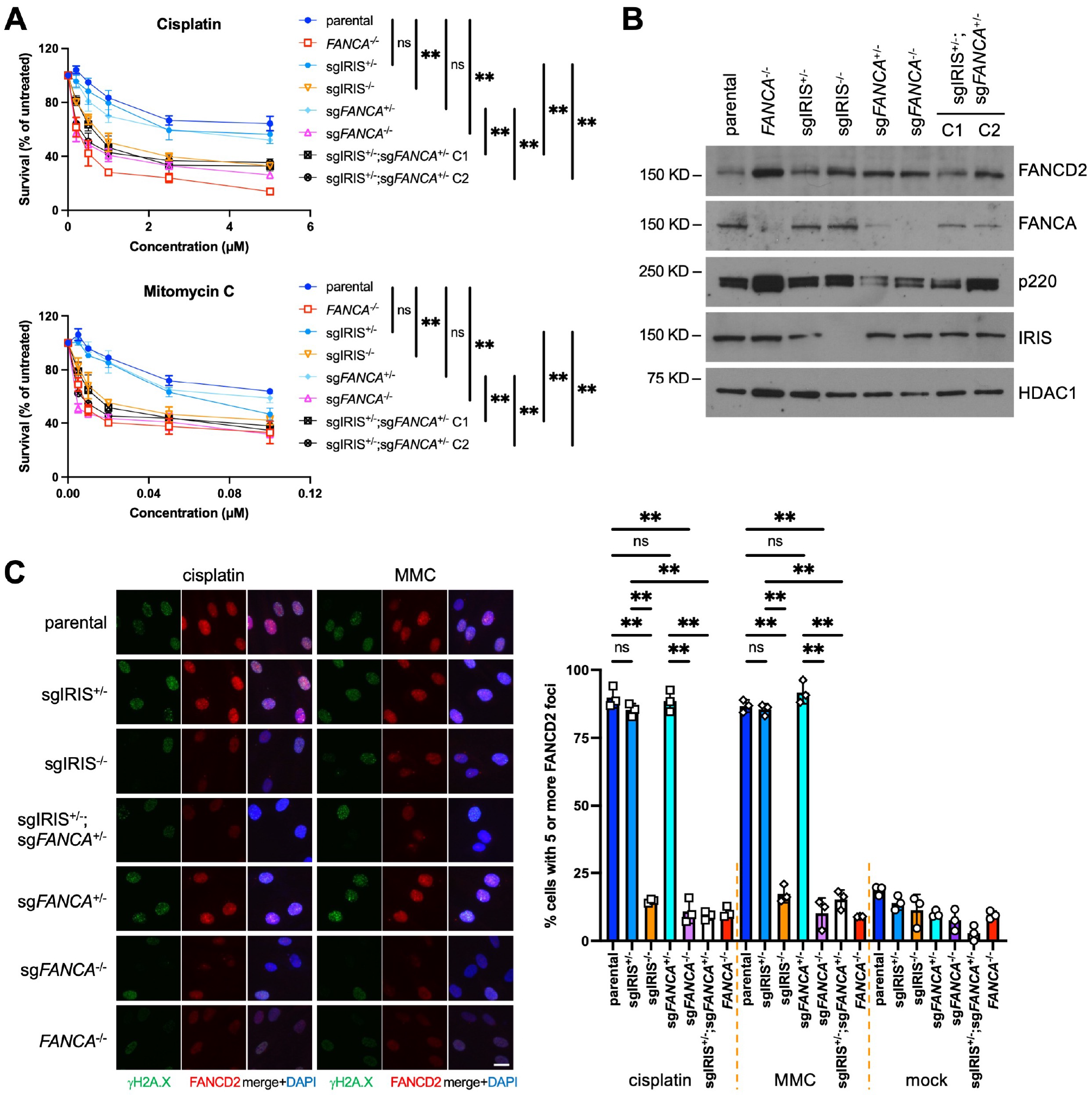
IRIS cooperates with FANCA in response to ICL damage. (A) Cooperation between IRIS and FANCA in cell survival after cisplatin or mitomycin C treatment. The IRIS and/or *FANCA* genotypes of the engineered fibroblasts are indicated. Fibroblasts were exposed to the indicated drug for 24 hours and growth was assayed with CellTiter-Glo after a total of 5 days. Data shown are normalized mean ± SD (n = 6) of two independent experiments with triplicate wells in each experiment. *P* values were obtained using a two-way ANOVA test. (B) Abundance of relevant proteins in the engineered fibroblasts analyzed by immunoblotting. (C) Analysis of ICL-induced FANCD2 foci in the engineered fibroblasts. Representative images of immunofluorescence of cells treated with the indicated drug for 24 hours before anti-γH2A.X and anti-FANCD2 immunostaining are shown. Quantifications of percentage of FANCD2 foci-positive cells are shown on the right. Bars represent mean ± SD (n = 3) of three independent experiments. *P* values were obtained using a two-tailed Student’s t test. Scale bar, 20 μm. ns, not significant; **p < 0.01.

We next analyzed ICL-induced FANCD2 foci in these engineered fibroblasts. In the compound heterozygote cells, although FANCD2 protein levels were comparable to those in parental cells (Figure 7B), FANCD2 was unable to form foci after cisplatin or MMC treatment (Figure 7C). These data suggest that IRIS operates with FANCA in the FA pathway to promote recruitment of FANCD2 to sites of ICL damage. Taken together, our data presented in this report strongly suggest that IRIS participates in ICL repair as an FA pathway protein, functioning upstream of the p220-directed HR, and that the association of IRIS with FANCA and FANCD2 plays an important role in ICL repair.

## Discussion

### Non-redundant functions of IRIS and p220 in the FA pathway

The canonical full-length protein product of the *BRCA1* gene, p220, has been characterized as FANCS in the FA pathway^3,15^. However, it was unclear why reconstitution of p220 expression was not sufficient for a full rescue of ICL-induced defects in *Brca1*-knockout cells^16,17^. In addition to p220, IRIS was deleted by the disruption of the *Brca1* gene in the knockout cells^16,17^. Thus, the partial rescue of ICL-induced phenotypes by p220 expression alone in *Brca1*-knockout cells suggests an incomplete restoration of ICL repair because of the absence of IRIS expression. Here, our findings reveal that IRIS is indeed required for ICL repair. IRIS operates upstream of the p220-directed HR in the FA pathway to promote FANCA and FANCD2 foci in response to ICL damage. In addition, IRIS is important, whereas p220 is dispensable, for ICL-inducible FANCD2 mono-ubiquitylation. The unique, *BRCA1* intron 11-encoded C-terminal tail of IRIS is important for both FANCD2 mono-ubiquitylation and recruitment to ICL sites. Moreover, although IRIS is not directly involved in HR, IRIS function is required for the p220-directed HR in the FA pathway. Hence, IRIS has critical functions that are unique from p220 function in ICL repair. Despite the extensive similarity in amino acid sequences between IRIS and p220, these *BRCA1* isoforms function non-redundantly in the FA pathway.

In addition to ICLs, p220 participates in repair of other types of DNA damage, including DSBs and SRF-associated damage^1,2,33^. Other FANC proteins, including FANCA and FANCD2, have also been shown to promote repair of these additional types of DNA damage^34-38^. By contrast with these FANC proteins, IRIS is neither required for cell survival after etoposide or HU treatment nor important for HR-mediated DSB repair. Hence, distinct from the ubiquitous requirement for p220 in DNA damage response and repair, IRIS function is restricted to the response and repair of ICLs.

### Distinct mechanisms by p220 and IRIS to promote FANCD2 recruitment to ICL sites

A possible mechanism for the function of p220 to promote ICL-inducible FANCD2 foci is the reported activity of p220 in unloading the CMG helicase from SRFs at ICL sites^14^ and thereby allowing downstream signaling. Our data argue that IRIS promotes ICL-inducible FANCD2 foci through a distinct mechanism. Following ICL induction, IRIS forms a protein complex with FANCA and FANCD2, which is independent of p220. The unique C-terminal tail of IRIS, which is not present on the p220 protein, is required for the complex formation and for FANCA and FANCD2 foci. We propose that IRIS promotes recruitment of FANCD2 to sites of ICL damage by connecting FANCD2 to the FA core complex at damage sites via IRIS-FANCA-FANCD2 complex formation.

Our data showed that p220 is not required for ICL-inducible FANCA foci (Figure 6A). Previously, p220 was implied to promote MMC-induced FANCA foci^39^. Of note, the siBRCA1 reagent assayed in the previous study targeted a common sequence of p220 and IRIS mRNAs^39^, suggesting that the observed defect in FANCA foci could be a result of a concurrent depletion of IRIS expression. In contrast, the shp220 and shIRIS reagents employed in the present study specifically targeted p220 and IRIS, respectively, without affecting expression of the other isoform (Figure S1A).

### Concomitant monoallelic mutations of different FA genes

FA is generally considered a recessive genetic disorder with biallelic mutations in one of at least 22 FA genes identified so far, except for patients containing X-linked *FANCB* or dominant-negative heterozygous *RAD51* mutations^7,8,27,28^. However, our data showed that cells with concomitant monoallelic mutations in IRIS and *FANCA* manifested defects in response to ICL damage, which phenocopied cells with biallelic mutations in *FANCA* (Figure 7). A similar observation that two FANC proteins cooperate in response to ICL damage has been reported. MEFs with hemizygous *Brca2*/*Fancd1* and hemizygous *Palb2*/*Fancn* (*Brca2*^KO/+^;*Palb2*^KO/+^) displayed a significant increase in chromosomal aberrations after MMC treatment when compared with *Palb2*^KO/+^ MEFs^40^.

Despite being extremely rare, diagnoses of FA patients with concomitant monoallelic mutations of different FA genes have been documented. For example, whole exome sequencing (WES) revealed that a patient inherited maternally a pathogenic mutation in *FANCA* and paternally a pathogenic mutation in *BRCA2*/*FANCD1*^41^. In another report, a patient carried a *de novo* monoallelic nonsense mutation in *BRCA2*/*FANCD1* plus an inherited heterozygous deletion in *PALB2*/*FANCN*, with no other sequence variants of known FA pathway genes being captured by WES^42^. In either case, however, the molecular diagnosis of the patient remained to be fully determined since WES could miss intronic variants of FA genes, some of which have been identified as pathogenic in FA patients^43^. Nonetheless, our data, along with these reports, suggest that FA can result from concomitant monoallelic mutations of two FA genes.

### Implication of the *BRCA1* intron 11-encoded IRIS tail in cancer

The *BRCA1* intron 11-encoded C-terminal tail unique to IRIS plays an important role in IRIS function in the FA pathway. Chronic, unrepaired ICL damage has been linked to tumor susceptibility, and germline mutations of FA genes have been implicated in increased risk of multiple cancers^6,8^. *BRCA1* variants in the intron 11-encoding sequence of IRIS have been identified (brcaexchange.org). One variant at the third nucleotide in the intron 11 sequence, named c.4096+3A>G, was suggested to display characteristics of a pathogenic mutation in hereditary breast and ovarian cancers^44^. This variant cosegregated with breast and ovarian cancer in several Icelandic carrier families. In tumor samples from heterozygous carriers in these families, a high incidence of loss of the wt *BRCA1* allele was observed^44^.

In addition to compromising the intron 11 donor splice site predicted in in silico analysis^45^, c.4096+3A>G leads to the replacement of isoleucine as the first amino acid residue of the C-terminal tail of IRIS protein with valine (Ile1367Val). The effect of this amino acid change on IRIS function in ICL repair is currently unclear. This and other variants in the intron 11-encoding sequence of IRIS, many of which result in missense and nonsense mutations in the unique C-terminal tail of IRIS, are understudied. Given our findings showing that the C-terminal tail is essential for IRIS function in ICL repair, it is necessary to study the effects of these mutations on cellular response and repair of ICLs and to examine their role in cancer development. A better understanding of the molecular function and potential pathogenic propensities of these IRIS variants will be beneficial for exploration of therapeutic interventions in cancer patients expressing such IRIS proteins.

## Materials and Methods

### Cells and Reagents

Maintenance of hTERT-immortalized human mammary epithelial (HME) cells was described previously^46^. Human skin fibroblasts, PD846F and RA3087, were maintained as described previously^47,48^. 293T cells were acquired from ATCC and grown in DMEM (Mediatech) plus 10% FBS (Invitrogen). All cells were verified to be mycoplasma-free.

Etoposide, hydroxyurea (HU), and mitomycin C (MMC) were purchased from Sigma Aldrich. Cisplatin was purchased from EMD Millipore.

### Cell Cycle Analysis

Analysis of cell cycle distribution was carried out as described previously^49^. Bromodeoxyuridine (BrdU) and the FITC Mouse Anti-BrdU Set were purchased from BD Biosciences. The Propidium Iodide (PI)/RNase Staining Solution was purchased from Cell Signaling. Cells were exposed to cisplatin or MMC for 24 hours prior to the BrdU pulse. Under each experimental condition, cells from three separate replicates of each culture were collected on an LSR Fortessa instrument (BD Biosciences) using the FACSDiva software (BD Biosciences) for data capture. The FlowJo (FlowJo, LLC) software was used for data analysis. All experiments were performed at least twice.

### Cell Survival/Growth Assays

Cells were exposed to different DNA-damaging agents 24 hours after seeding in triplicate wells (2000 cells/well) for each drug concentration in 96-well plates. Drugs were washed away after 24-hour incubation. Five days later, cell growth was assayed with CellTiter-Glo (Promega) following the manufacturer’s protocol. All experiments were performed twice.

### Comet Assay

Alkaline comet assays were performed using the Single-Cell Gel Electrophoresis Assay kit (4250-050-K, Trevigen) according to the manufacturer’s instruction. All experiments were performed twice. The CellProfiler software was used for data analysis and quantitation. A total of 100-200 nucleoids per condition was analyzed.

### CRISPR Editing

Synthesized IRIS guide RNA oligonucleotide (target sequence: 5’-CTGGGGCAAACACAAAAACCTGG-3’) was cloned into the BsmB1 sites of the lentiCRISPRv2 vector (a gift from Feng Zhang, Addgene plasmid #52961)^50^ following the protocol described by Sanjana et al.^50^. FANCA guide RNA oligonucleotides (target sequences: 5’-GCGCCTCCTGCGAAGCCATCAGG-3’ and 5’-CGTAGCGGGAAGGGTCAAGAGGG-3’) were similarly cloned into the lentiCRISPRv2-hygro vector (a gift from Brett Stringer, Addgene plasmid #98291)^51^.

Generation of engineered PD846F fibroblasts was performed in a similar way as described previously^52^. Briefly, sgRNA vectors were transfected into PD846F fibroblasts using Lipofectamine 2000 (Life Technologies). Transfected cells were selected by puromycin or hygromycin (Roche) for 4 days, after which cells were cultivated for 14 more days in antibiotic-free medium before colony-picking. Genetic statuses of corresponding genes in clonal cells were verified by sequencing.

### ICL and DSB Repair Assays

Participations of p220 and IRIS in ICL and DSB repair were analyzed using an ICL repair reporter assay and a DSB repair reporter assay, respectively^4,22^. Crosslinking and control oligonucleotides were synthesized by Integrated DNA Technologies. Cell preparation for fluorescence-activated cell sorting (FACS) analysis was performed essentially as described previously^4,22^. Cells from three separate replicates of each culture under each assay condition were analyzed on an LSR Fortessa instrument (BD Biosciences) using the FACSDiva software (BD Biosciences) for data capture. All experiments were performed at least twice. The FlowJo (FlowJo, LLC) software was used for data analysis.

### Antibodies, Immunoblotting, Immunofluorescence, and Immunoprecipitation (IP)

The following antibodies were used: γH2A.X (05-636-I, EMD Millipore; #9718, Cell Signaling), FANCD2 (NB100-182, Novus), FANCA (NBP2-56898, Novus; #14657, Cell Signaling), FANCL (PA5-57768, Invitrogen), phospho-CHK1 (#2348, Cell Signaling), HDAC1 (#5356, Cell Signaling), Histone H3 (#4499, Cell Signaling), BRCA1 (A300-000A, Bethyl; OP92, EMD Millipore; sc-6954, Santa Cruz), Actin (A5441, Sigma), HA (901501, BioLegend), Myc (sc-40, Santa Cruz), and V5 (A190-120A, Bethyl). Anti-human IRIS antibody was described previously^46^.

Western blotting and IP were carried out as described previously^53^ with specific antibodies. Nuclear soluble and chromatin fractions were prepared and combined as nuclear extracts (NEs) using a Subcellular Protein Fractionation Kit for Cultured Cells (Thermo Scientific). All experiments were performed at least twice.

Immunofluorescence analyses were performed as described previously^54^. A total of 100-200 cells per condition from 3 independent experiments was analyzed for foci determination. Images in Figures 4A, 4B, and S4B were acquired using a Zeiss LSM 980 confocal microscope controlled by Zen software. Image analysis was performed in FIJI.

### Plasmids and Mutagenesis

The wt and Tr IRIS constructs were described previously^46^. The FANCA and FANCD2 expression vectors were gifts from Jacob Corn (Addgene plasmids #111126 and #111127)^55^. The FANCD2 ΔTower construct was generated using a QuikChange Site-Directed Mutagenesis kit (Agilent) according to the manufacturer’s protocol. The FANCL-V5 construct was generated by cloning FANCL cDNA into the pLX304 vector through Gateway cloning (Life Technologies). All constructs were confirmed as accurate by sequencing.

### siRNA and shRNA Experiments

Transfection of siRNA was performed using Lipofectamine 2000 (Life Technologies) according to the manufacturer’s protocol. Target sequences of siRNA were described previously^46,56^. Target sequences of shRNA and generation of cells carrying shRNA were described previously^46,54^.

### Statistical Analysis

Data are presented as *mean* ± *SD*, except when indicated otherwise. The *P* values were calculated using a two-way ANOVA test, a two-tailed Student’s t test, or a Wilcoxon rank-sum test (two-tailed) as indicated. The Holm-Bonferroni adjustment was used for multiple comparisons. *P* < 0.05 is considered significant.

## Data Availability

The authors declare that all the data supporting the findings of this study are available within the article and its supplemental information file, and from the corresponding authors upon reasonable request.

## Acknowledgments

We are grateful to Y. Li for technical assistance. We are in debt to M. Jasin for providing the ICL repair reporter system. We are thankful to A. Smogorzewska for sharing RA3087 fibroblasts. We are thankful to F. Zhang, B. Stringer, and J. Corn for providing the CRISPR, FANCA, and FANCD2 plasmids. We thank T. Roberts and R. Weinberg for critical reading of the manuscript. We thank the Livingston laboratory for helpful discussions and D. Livingston for his endless inspiration and encouragement. We thank the Molecular Imaging Core of Dana-Farber Cancer Institute for the use of microscopes and support in imaging.

This body of work was supported by grants to D.M.L. from NCI (5P01CA080111), from the DFCI/Novartis Program in Drug Discovery, from the Breast Cancer Research Foundation (BCRF-21-101, BCRF-20-101, BCRF-19-101, and BCRF-18-101), and from the Gray Foundation.

## Author Contributions

A.G.L., S.B.C., J.S.B., M.B., and D.M.L. designed the study. A.G.L., B.C.C., E.C.M., Y.H., M.O., and Q.K. performed the experiments. A.G.L., B.C.C., E.C.M., and D.M.L. analyzed the data. A.G.L., S.B.C., J.S.B., and M.B. wrote the manuscript.

