## Supplemental Figures for "The *BRCA1* isoform, BRCA1-IRIS, operates independently of the full-length BRCA1 in the Fanconi anemia pathway"

### Figure S1. IRIS function in the cellular response to DNA damage is restricted to ICLs

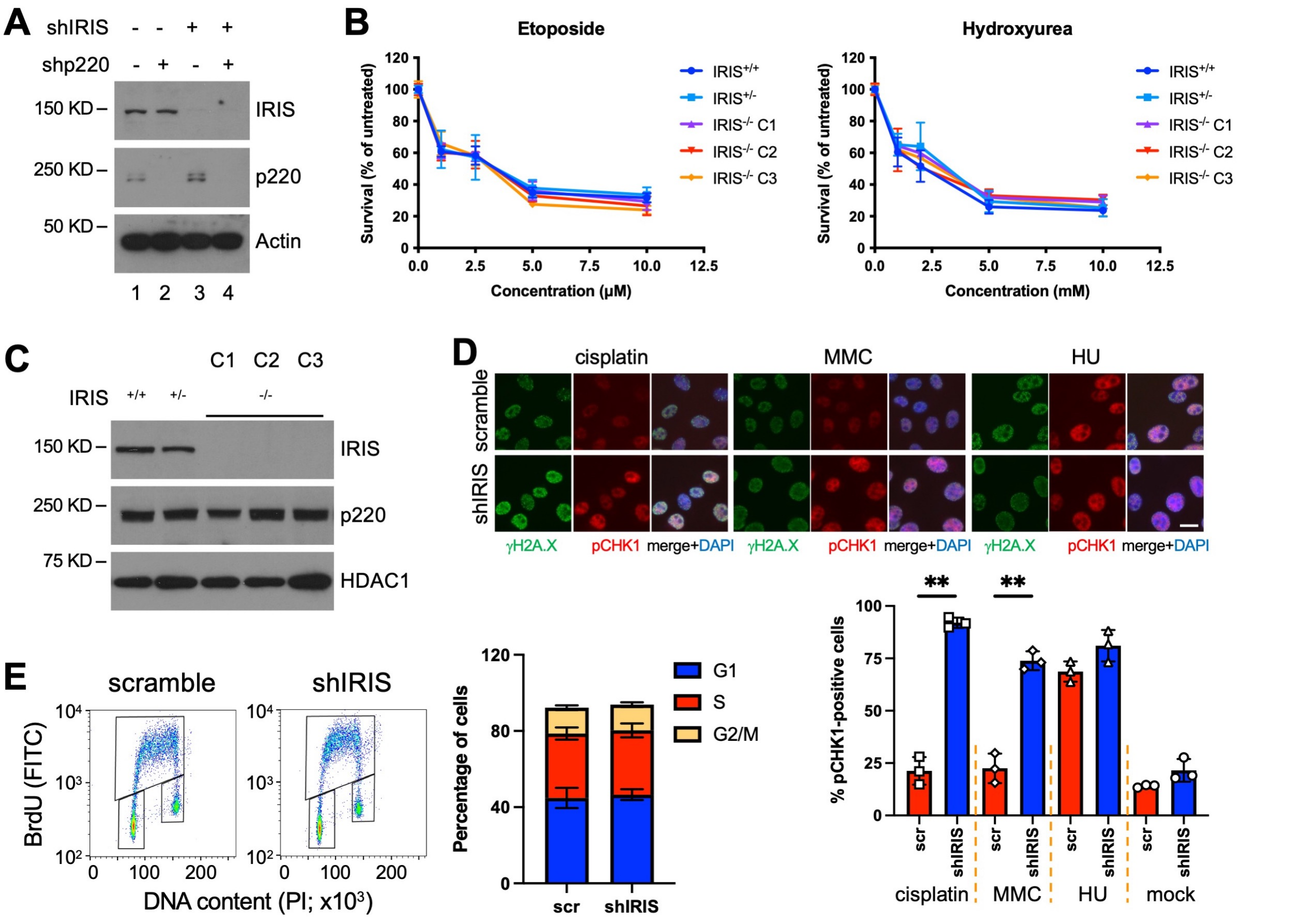


**Figure S1. IRIS function in the cellular response to DNA damage is restricted to ICLs, related to Figure 1.**

(A) Western blots showing IRIS and p220 protein levels in HME cells expressing various combinations of the indicated hairpins.

(B) Survival of IRIS-knockout fibroblasts in response to etoposide or hydroxyurea. Fibroblasts with the indicated IRIS genotype were exposed to etoposide or hydroxyurea for 24 hours and growth was assayed with CellTiter-Glo after a total of 5 days. Data shown are normalized mean ± SD (n = 6) of two independent experiments with triplicate wells in each experiment.

(C) Western blots showing IRIS and p220 protein levels in the indicated fibroblasts.

(D) Exposure of IRIS-depleted HME cells to DNA interstrand crosslinkers led to appearance of phospho-CHK1 signal in nuclei. Representative images of immunofluorescence of cells treated with the indicated drug for 24 hours before anti-γH2A.X and anti-pCHK1$^{S345}$ immunostaining are shown. Quantifications of percentage of cells positive for phospho-CHK1 signal under each treatment condition are shown on the bottom. Bars represent mean ± SD (n = 3) of three independent experiments. P values were obtained using a two-tailed Student's t test. **p < 0.01. MMC, mitomycin C; HU, hydroxyurea; scr, scramble. Scale bar, 20 μm.

(E) IRIS depletion had no effect on cell cycle distribution. Representative plots of fluorescence-activated cell sorting (FACS) analysis are shown. Summaries of mean percentage of cells ± SD (n = 6) in each cell cycle phase of two independent experiments with triplicate cultures in each experiment are shown on the right.

### Figure S2. Effects of IRIS expression on FANCD2 or UHRF1 foci and cell cycle

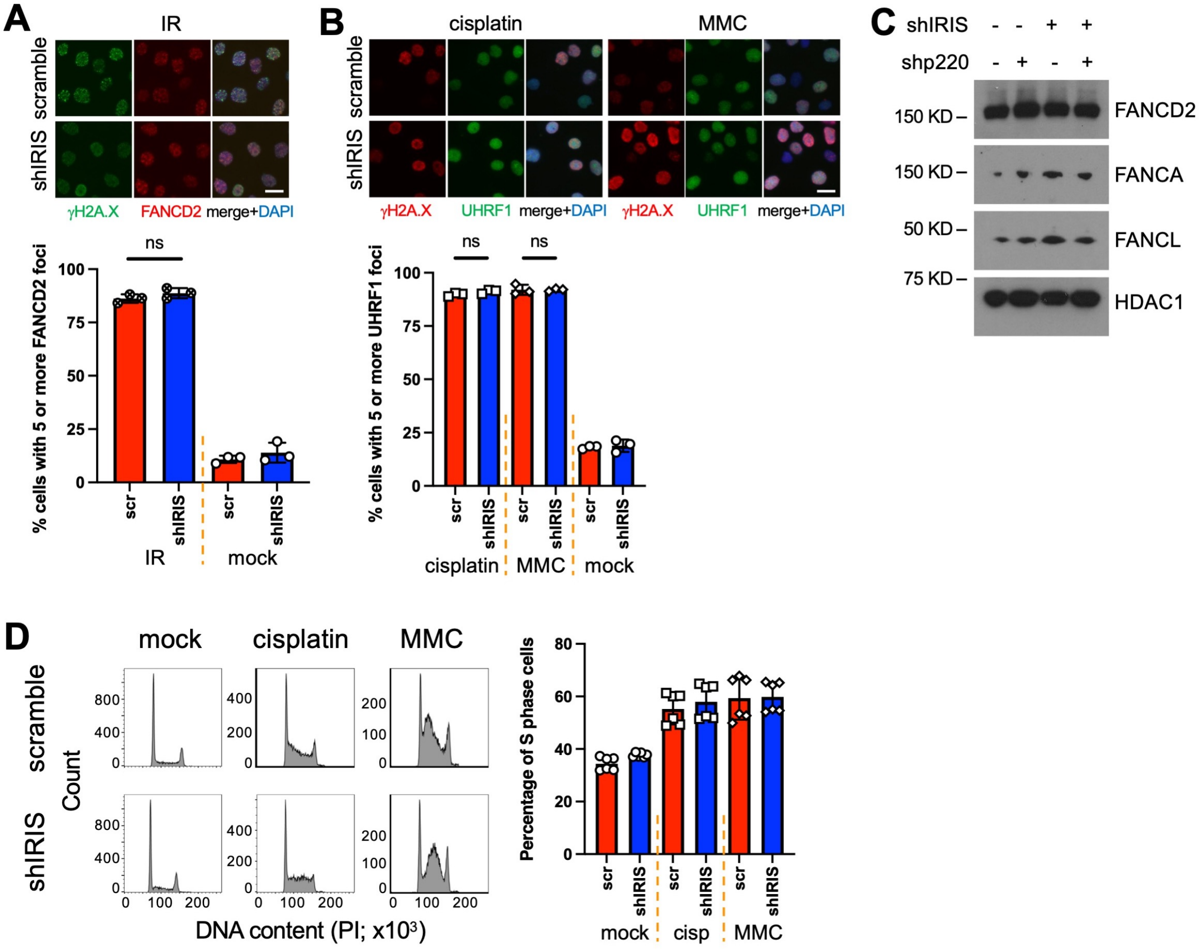


**Figure S2. Effects of IRIS expression on FANCD2 or UHRF1 foci and cell cycle, related to Figure 2.**

(A) γ-irradiation (IR) induced FANCD2 foci in both control and IRIS-depleted cells. Quantifications of percentage of FANCD2 foci-positive cells are shown on the bottom.

(B) ICL-induced UHRF1 foci were not dependent on IRIS. Quantifications of percentage of UHRF1 foci-positive cells are shown on the bottom.

(C) Expression levels of certain FANC proteins in HME cells examined by immunoblotting.

(D) Cell cycle analysis of IRIS-depleted HME cells exposed to DNA interstrand crosslinkers. Representative FACS histograms are shown. Bars representing mean percentage ± SD (n = 6) of S phase cells of two independent experiments with triplicate cultures in each experiment are shown on the right. cisp, cisplatin.

Representative images of immunofluorescence of HME cells treated with the indicated drug for 24 hours or incubated for 4 hours post IR (10 Gy) before immunostaining using the indicated antibodies are shown. Bars, except in (D), represent mean ± SD (n = 3) of three independent experiments. P values were obtained using a two-tailed Student's t test. ns, not significant. Scale bar, 20 μm.

### Figure S3. Damage type-specific formation of the IRIS-FANCA-FANCD2 complex

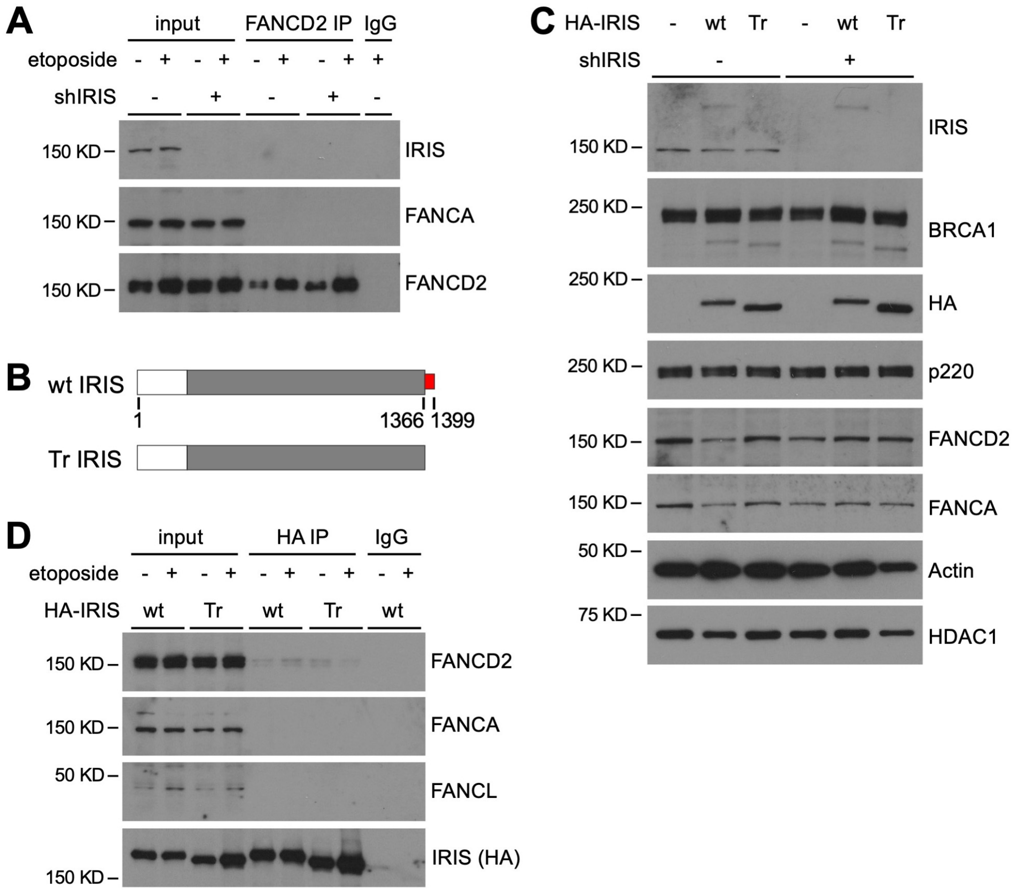


**Figure S3. Damage type-specific formation of the IRIS-FANCA-FANCD2 complex, related to Figure 3.**

(A) No association between endogenous IRIS and FANCD2 in HME cells exposed to etoposide. Nuclear extracts (NEs) of the indicated HME cells were prepared after 24-hour etoposide or mock treatment and used for IP with FANCD2 antibody followed by immunoblotting.

(B) A schematic representation of the wild-type (wt) IRIS protein and a truncated (Tr) mutant IRIS protein used in this study. Numbers indicate positions of amino acid residues of the wt protein. Tr IRIS lacks the amino acid residues encoded by BRCA1 intron 11.

(C) Immunoblots showing expression levels of different IRIS species (wt, Tr, and endogenous) and certain FANC proteins in control and IRIS-depleted HME cells.

(D) Etoposide did not stimulate binding of certain FANC proteins to wt or Tr IRIS. After 24-hour drug treatment, NEs from the indicated HME cells were used for IP with HA antibody followed by Western blotting.

### Figure S4. The C-terminal tail of IRIS is not required for etoposide-induced FANCD2 foci

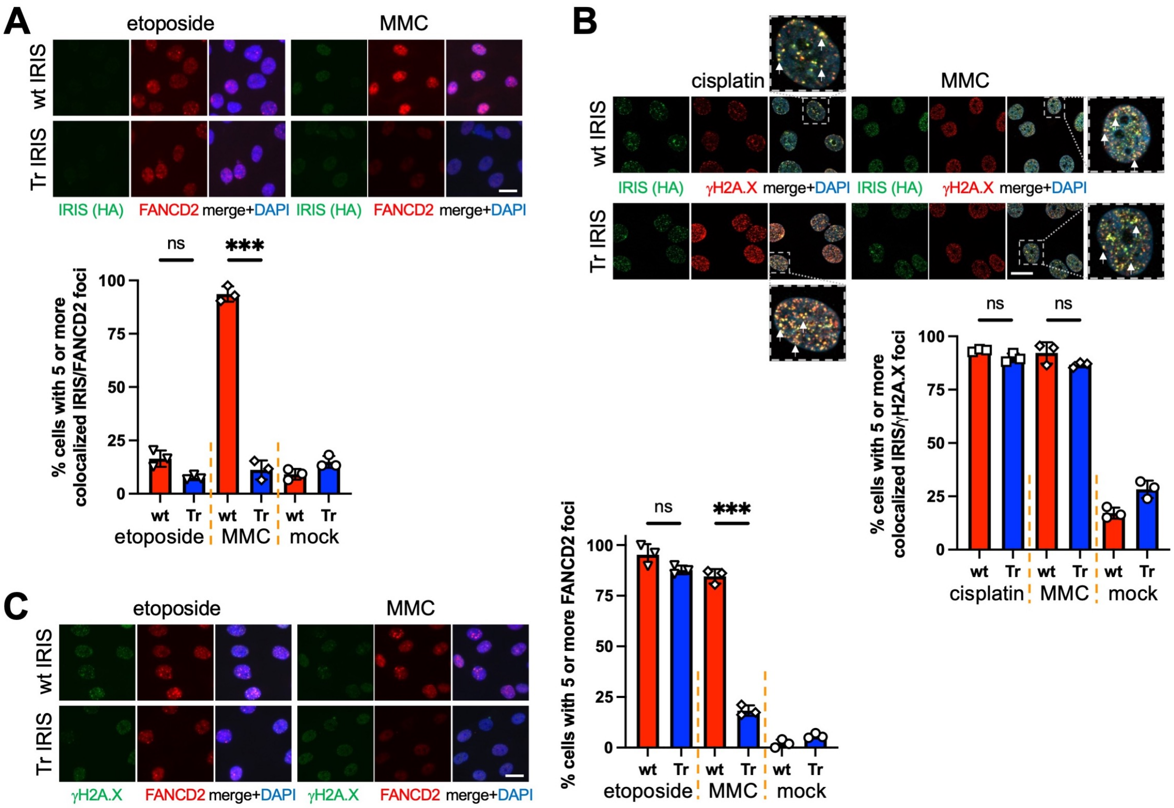


**Figure S4. The C-terminal tail of IRIS is not required for etoposide-induced FANCD2 foci, related to Figure 4.**

(A) Neither wt nor Tr IRIS appeared in foci after etoposide treatment. Quantifications of percentage of cells positive for IRIS/FANCD2-overlapping foci are shown on the bottom.

(B) Both wt and Tr IRIS appeared in γH2A.X-containing foci after cisplatin or MMC treatment. Arrows point to some IRIS/γH2A.X colocalized foci. Zoomed-in areas containing a single nucleus each are shown next to their corresponding merged panels. Quantifications of percentage of IRIS foci-positive cells are shown on the bottom.

(C) FANCD2 foci were detected in both wt and Tr IRIS-expressing HME cells exposed to etoposide. Quantifications of percentage of FANCD2 foci-positive cells are shown on the right.

Representative images of immunofluorescence of HME cells treated with the indicated drug for 24 hours before immunostaining using the indicated antibodies are shown. Bars represent mean ± SD (n = 3) of three independent experiments. P values were obtained using a two-tailed Student's t test. ns, not significant; ***$p < 0.001$. Scale bar, 20 μm.

### Figure S5. Effects of IRIS expression on etoposide-induced FANCD2 mono-ubiquitylation and cell survival after drug treatment

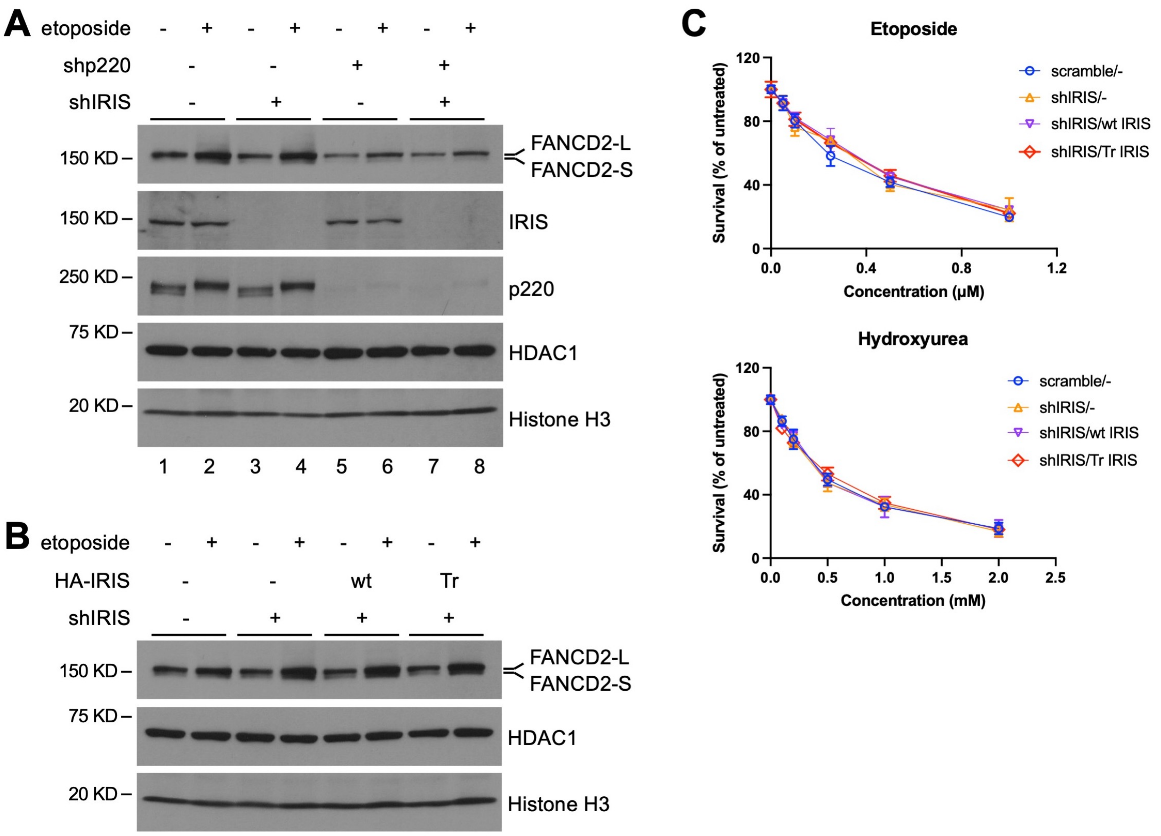


**Figure S5. Effects of IRIS expression on etoposide-induced FANCD2 mono-ubiquitylation and cell survival after drug treatment, related to Figure 5.**

(A) IRIS was not important for etoposide-induced FANCD2 mono-ubiquitylation. NEs of the indicated HME cells were prepared after 24-hour drug treatment and analyzed by immunoblotting.

(B) Neither wt nor Tr IRIS expression affected mono-ubiquitylation of FANCD2 in HME cells exposed to etoposide. After 24-hour drug treatment, NEs from the indicated HME cells were used for Western blotting analysis.

(C) Expression of neither wt nor Tr IRIS in IRIS-depleted HME cells influenced cell survival in response to etoposide or hydroxyurea. HME cells expressing the indicated hairpin and cDNA were exposed to etoposide or hydroxyurea for 24 hours and growth was assayed with CellTiter-Glo after a total of 5 days. Data shown are normalized mean ± SD (n = 6) of two independent experiments with triplicate wells in each experiment.

### Figure S6. Effects of IRIS expression on FANCD2 or p220 foci

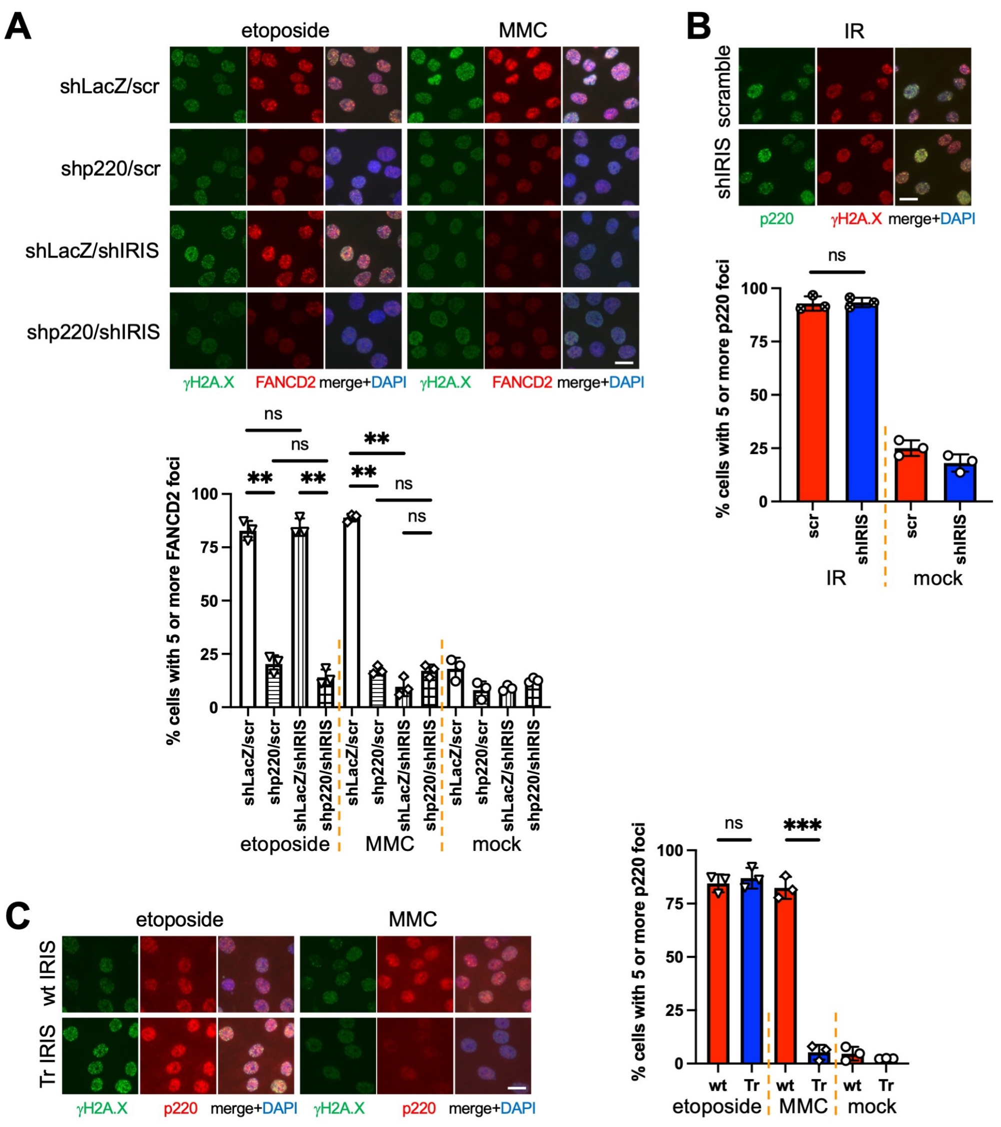


**Figure S6. Effects of IRIS expression on FANCD2 or p220 foci, related to Figure 6.**

(A) FANCD2 foci depended on both p220 and IRIS expression in MMC-treated HME cells, while its foci were affected by loss of p220, but not IRIS, in etoposide-treated cells. Quantifications of percentage of FANCD2 foci-positive cells are shown on the bottom.

(B) In response to γ-irradiation (IR), p220 foci appeared in both IRIS-depleted and control cells. Quantifications of percentage of cells positive for p220 foci are shown on the bottom.

(C) Etoposide-induced p220 foci were not regulated by expression of wt or Tr IRIS. Quantifications of percentage of p220 foci-positive cells are shown on the right.

Representative images of immunofluorescence of HME cells treated with the indicated drug for 24 hours or incubated for 4 hours post IR (10 Gy) before immunostaining using the indicated antibodies are shown. Bars represent mean ± SD (n = 3) of three independent experiments. P values were obtained using a two-tailed Student's t test. ns, not significant; $**p < 0.01$; $***p < 0.001$. Scale bar, 20 μm.
